# Topoisomerase III-ß is required for efficient replication of positive-sense RNA viruses

**DOI:** 10.1101/2020.03.24.005900

**Authors:** K. Reddisiva Prasanth, Minato Hirano, W. Samuel Fagg, Eileen T. McAnarney, Chao Shan, Xuping Xie, Adam Hage, Colette A. Pietzsch, Alexander Bukreyev, Ricardo Rajsbaum, Pei-Yong Shi, Mark T. Bedford, Shelton S. Bradrick, Vineet Menachery, Mariano A. Garcia-Blanco

**Author notes:** Correspondence to: Mariano A. Garcia-Blanco. Global Health, Surveillance & Diagnostics Division, MRIGlobal, 425 Volker Blvd, Kansas City, MO 64110.

## Abstract

Based on genome-scale loss-of-function screens we discovered that Topoisomerase III-ß (TOP3B), a human topoisomerase that acts on DNA and RNA, is required for yellow fever virus and dengue virus-2 replication. Remarkably, we found that TOP3B is required for efficient replication of all positive-sense-single stranded RNA viruses tested, including SARS-CoV-2. While there are no drugs that specifically inhibit this topoisomerase, we posit that TOP3B is an attractive anti-viral target.

---

Acute viral infections, particularly those caused by RNA viruses, are dangerous threats to public health. This has been made evident by the ongoing pandemic of severe acute respiratory syndrome (SARS), officially named COVID-19, caused by the betacoronavirus SARS-CoV-2. When confronted with emerging viruses such as SARS-CoV-2 we are left with few countermeasures that can be deployed to specifically prevent or treat the diseases they cause. We and others have argued that deeper understanding of the molecular mechanisms required for viral replication can reveal potential vulnerabilities for related viruses. These vulnerabilities could be based on unique viral targets, such as the RNA-dependent RNA polymerases, or on dependency on host encoded pro-viral activities. To identify such host pro-viral factors we embarked on genome-scale screens for diverse families of viruses and identified scores of required host factors ^1–4^. Some of these host factors are attractive drug targets.

A meta-analysis of RNAi-based loss-of-function screens for YFV and DENV-2 host factors revealed 274 common candidates ^1^. TDRD3, a Tudor domain containing protein that interacts with methylated arginine motifs ^5^, was identified as a candidate host factor required for both DENV2/YFV. In the two YFV screens TDRD3 ranked 66^th^ of over 21,500 gene products in terms of how well its knockdown decreased YFV (adjusted p value = 0.0006). CRISPR-Cas9 mediated knockout of TDRD3 in HuH-7 cells confirmed that this protein was required for efficient DENV-2 replication (Fig 1A & B).

**Figure 1.**
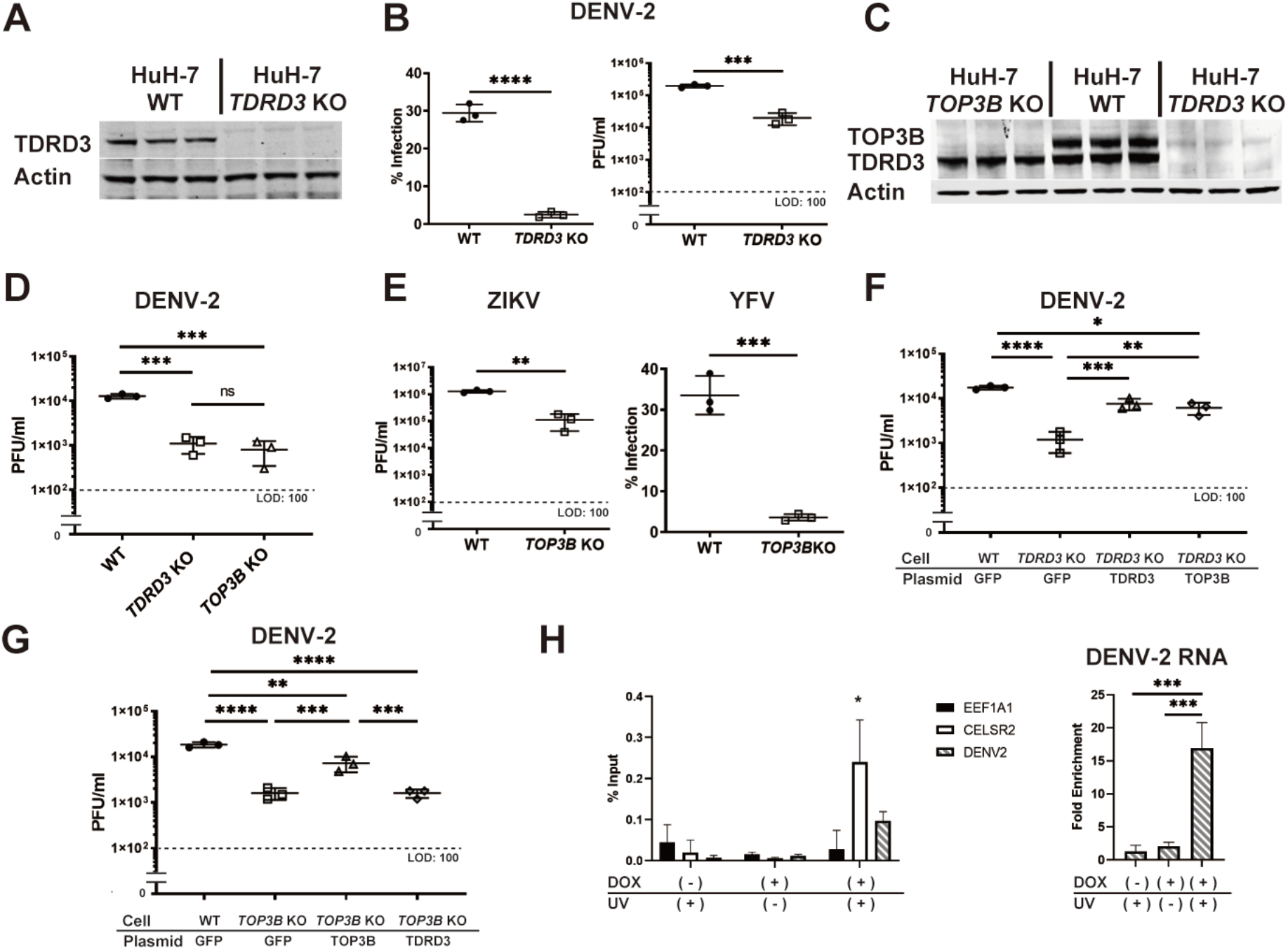
TOP3B is required for efficient replication of multiple flaviviruses. (**A**) TDRD3 expression in HuH-7 and TDRD3 KO cells. (**B**) TDRD3 KO inhibits DENV-2 infectivity (left) and propagation (right) (**C**) TOP3B and TDRD3 expresssion in HuH-7, TDRD3 KO, and TOP3B KO cells. (**D**) TOP3B KO inhibits DENV-2 propagation. (**E**) TOP3B KO inhibits ZIKV (left) and YFV-17D (right) propagation. (**F**) TOP3B overexpression rescues TDRD3 KO. (**G**) TDRD3 overexpression does not rescue TOP3B KO. (**H**) TOP3B can be crosslinked to DENV-2 RNA during infection. *: p < 0.05, **: p < 0.01, ***: p<0.001 and ****: p < 0.0001

A well-known function of TDRD3 is to bind and stabilize Topoisomerase III-ß (TOP3B) ^6–8^, a type IA topoisomerase and the only human topoisomerase known to act on both DNA and RNA ^6,9^. Knockout of TDRD3 in HuH-7 cells led to levels of TOP3B that were almost as low as those obtained with knockout of TOP3B itself (Fig 1C). Knockout of TOP3B, which does not alter TDRD3 levels, resulted in the same dramatic decrease in DENV-2, YFV and Zika virus (ZIKV) replication as knockout of TDRD3 (Fig 1D & E), which suggested that the only role of TDRD3 in viral replication was to stabilize TOP3B. Indeed, TOP3B overexpression rescued DENV-2 infection in TDRD3 KO cells (Fig 1F). The reverse was not true, TDRD3 overexpression was incapable of rescuing virus replication in a TOP3B KO cells. Therefore, we conclude that TOP3B is a proviral host factor for several flaviviruses.

The genetic approaches we utilized above do not distinguish between direct and indirect modes of action. To address whether or not TOP3B directly interacted with DENV-2 genomes we used UV crosslinking followed by RNA immunoprecipitation (CLIP). We carried out CLIP assays using a HEK-293T cell line that expressed a FLAG-tagged TOP3B upon doxycycline treatment and anti-FLAG antibodies to carry out the immunoprecipitation. FLAG-TOP3B preferentially crosslinked to CELSR2 RNA, which was previously known to bind this topoisomerase ^6^, relative to EEF1A1 RNA, which we use as a negative control (Fig 1H). Importantly, TOP3B crosslinked DENV-2 RNA (Fig 1H), strongly suggesting that TOP3B acts directly on the viral genome.

Since TOP3B was required for DENV-2, YFV and ZIKV replication, we asked whether or not this topoisomerase was required for replication of other viruses. Influenza A virus, which has a negative-sense segmented RNA genome and belongs to the family *Orthomyxoviridae*, replicated significantly better in TOP3B KO cells than in the parental HuH-7 cells (Fig 2A, left panel), suggesting that TOP3B plays an anti-viral role for influenza A virus. We also tested Ebola virus, a member of the family *Filoviridae*, which has a negative sense RNA genome and again noted that Ebola virus replicated slightly better in TOP3B KO cells than in the parental HuH-7 cells (Fig 2A, right panel). We thus suggest that TOP3B may be an anti-viral factor for negative sense RNA viruses, and clearly is not required for the replication of influenza A virus or Ebola virus.

**Figure 2.**
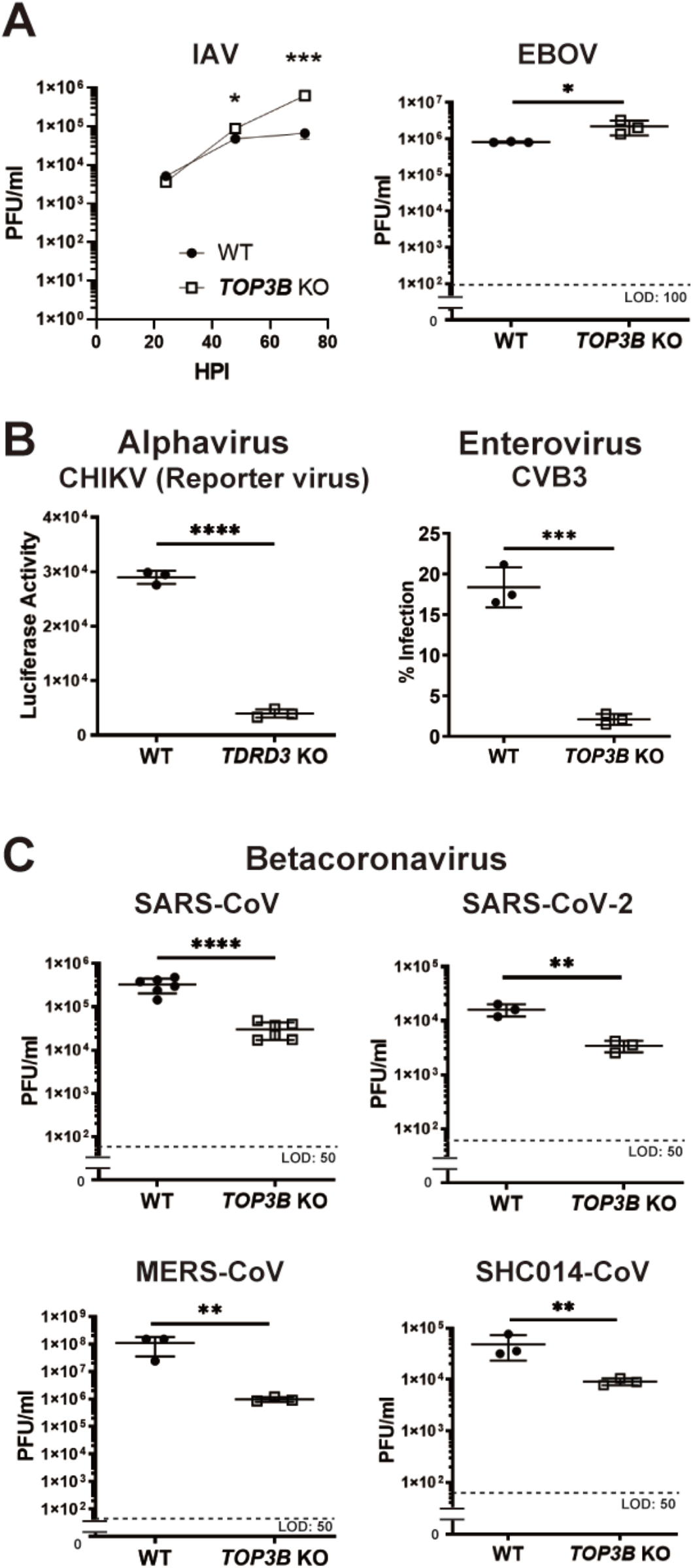
TOP3B is required for efficient replication of a diverse group of (+) ss RNA viruses. (**A**) Influenza A virus (IAV) (left) and Ebola virus (EBOV) (right) replicate better in TOP3B KO cells. (**B**) TDRD3 KO inhibits chikungunya virus replication (left) and TOP3B KO inhibits coxsackievirus B3 replication (right). (**C**) TOP3B KO inhibits replication of SARS-CoV, SARS-CoV-2, MERS-CoV, and SCH1014-CoV. *: p < 0.05, **: p < 0.01, ***: p<0.001 and ****: p < 0.0001

We tested a series of positive-sense single-stranded RNA viruses ((+) ss RNA viruses). A recombinant chikungunya virus (CHIKV), an alphavirus of the *Togaviridae* family was sensitive to TDRD3 knockout (Fig 2B, left panel), and coxsackievirus B3 (CVB3), an enterovirus of the family *Picornaviridae*, was dependent on TOP3B for efficient replication (Fig 2B, right panel). Most important in the context of the COVID-19 pandemic, four betacoronaviruses, of the *Coronaviridae* family, SARS-CoV, SARS-CoV-2, MERS-CoV, and SCH1014-CoV, a bat coronavirus, were significantly crippled by TOP3B KO (Fig 2C). These results indicated that TOP3B is a host factor essential for efficient replication of a diverse group of (+) ss RNA viruses.

Among host factors required for diverse groups of RNA viruses are components of the translation machinery and the proteasome, in addition to factors that mediate membrane transactions required for entry and exit of viruses. The understanding of the broad requirement for TOP3B among (+) ss RNA viruses will likely provide important insights into heretofore understudied steps in their lifecycles. While a requirement for human topoisomerase TOP1 in exacerbated inflammation induced by RNA virus infection has been previously documented ^10^, the direct action of an RNA topoisomerase in viral replication is unprecedented, and has important biological implications. A requirement for a RNA topoisomerase as a critical cofactor for (+) ss RNA viruses is reasonable since viral genomes, because of their complex structures and size, especially the large coronaviruses, may lend themselves to topological problems (knots and catenanes) ^11,12^. In parallel work focused on exploring these implications we have shown that TOP3B is required for a late step in DENV-2 replication, likely during the assembly of the genome into virions (Prasanth et al., manuscript in preparation). Furthermore, this action requires the topoisomerase activity of TOP3B since it is abrogated by a tyrosine to phenylalanine mutation (Y336F) in the active site of the enzyme (Ibid).

Here we emphasize the potential for TOP3B as a drug target. It must be clear that we make this suggestion responsibly with full understanding that our work is an early step in what must be a rigorous process to identify effective antivirals. Nonetheless, several facts support this consideration. TOP3B^-/-^ mice are viable ^13^ and humans with a homozygous 240-kb deletion of chromosome 22q11.22 spanning TOP3B gene are also viable^7^. Although both homozygous null mice and humans have abnormal phenotypes, the enzyme is not essential. Topoisomerase inhibitors are well known drugs used to treat cancer (e.g. doxorubicin)^14^ and bacterial infections (e.g., ciproflaxin) ^15^. Unfortunately, we are not aware of any approved TOP3B drugs and prior to this report there was no evidence that inhibiting TOP3B could be a useful in treating acute viral infections. In fact other than the ACE-2 receptor, and perhaps furin, required for SARS-CoV-2 attachment and entry ^16^, TOP3B is the only other known required host factor. We hope this brief communication will spur interactions between individuals studying TOP3B and developing inhibitors of TOP3B, and those working on SARS-CoV-2.

## Acknowledgements

We thank members of our laboratories at the University of Texas Medical Branch for comments and suggestions. We also thank Dr. Yuk-Ching Tse Dinh (Florida International University) and Dr. Yves Pommier (National Cancer Institute) and members of their laboratories, and Dr. Phillip A. Sharp (MIT) for important discussions and comments. We acknowledge support from the Uehara Foundation Fellowship (MH), NIH R01 CA204806, Vacek Chair, and UTMB startup package (MGB), NIH R01 GM126412 (MTB), NIH AI142759, and awards from the Kleberg Foundation, John S. Dunn Foundation, Amon G. Carter Foundation, Gilson Longenbaugh Foundation, and Summerfield Robert Foundation (PYS), NIH/NIAID T32 AI060549 (AH), R21 AI26012 and R01 AI134907 (RR), UTMB start up funds (VM).

## Author contributions

KRP did the majority of the experimental work in Figure 1 and coordinated the work in Figure 2, MH did the CLIP experiments in Figure 1, analyzed the data for all figures and prepared final figures. CS, XX and PYS made the reporter CHIKV and provided essential reagents for flavivirus work. MGB wrote the first draft of the manuscript, except the online methods, which was drafted by MH and KRP. MTB provided critical reagents and expertise needed to start the work on this project. WSF, SSB and MGB supervised the work in the Garcia-Blanco laboratory. AH performed and analyzed, and RR supervised work on IAV. CAP performed and AB supervised work with EBOV.

## Methods (to be included only online)

### Cells, and Viruses

Human hepatoma cells (HuH-7) and African green monkey kidney (Vero) cells were kindly provided by Charles Rice (Rockefeller University), and maintained in Dulbecco’s modified Eagle’s medium (DMEM) (Gibco) supplemented with 10% fetal bovine serum (FBS), 100 U/ml penicillin and 100 μg/ml streptomycin at 37°C with 5% CO_2_. Baby Hamster Kidney-21 (BHK21) cells were maintained in Roswell Park Memorial Institute medium (RPMI) (Gibco) supplemented with 10% FBS, 100 U/ml penicillin and 100 μg/ml streptomycin at 37°C with 5% CO_2_. Human Embryonic kidney 293 with FLP-In site (HEK293 Flp-In T-REx) cells were purchased from Thermo Fisher Scientific, and maintained in Dulbecco’s modified Eagle’s medium (DMEM) (Gibco) supplemented with 10% FBS, 100 U/ml penicillin, 100 μg/ml streptomycin, zeocin (100 μg/ml) and blasticidin (15 μg/ml) at 37°C with 5% CO_2_.

Dengue virus-2 (DENV-2 New Guinea C), yellow fever virus (YFV 17D) or Coxsackie virus B3 (CVB3 Strain 20) were prepared as working stock after the virus propagation in C6/36 (DENV-2, YFV) or HeLa (CVB3), respectively ^3^. Zika virus (ZIKV-Cambodian) was derived from infections cDNA clone and propagated in Vero cells as described before ^17^. A Renilla luciferase reporter chikungunya virus (CHIKV) was derived from an infectious cDNA clone where the luciferase gene was inserted into a subgenomic regions (Unpublished; Chao Shan, Xuping Xie, and Pei-Yong Shi). The recombinant Ebola virus (EBOV) strain Mayinga expressing green fluorescent protein (GFP) from the GFP gene inserted between the NP and VP35 genes was recovered from the EBOV full-length clone which were provided by Jonathan Towner and Stuart Nichol (Centers for Disease Control and Prevention [CDC]) and the support plasmids provided by Drs. Y. Kawaoka (University of Wisconsin) and H. Feldmann (Rocky Mountain Laboratories, NIAID) and was identical to the virus described previously ^18^. IAV (Influenza A/Puerto Rico/8/1934 H1N1) titers were determined by plaque assays. SARS-CoV, SARS-CoV-2, MERS-CoV, and SCH014-CoV were obtained from the The World Reference Center for Emerging Viruses and Arboviruses at the Galveston National Laboratory.

Experiment using infectious EBOV was conducted in the Bio-Safety Level (BSL) 4 facility at the Galveston National Laboratory, University of Texas Medical Branch. Experiments using SARS-CoV, SARS-CoV-2, MERS-CoV, and SCH014-CoV were conducted in BSL3 laboratories. Other viruses were handled in BSL2 conditions until the proper inactivation and disinfection.

### Virus titration

To determine virus titer, plaque-forming assays of DENV were performed by using BHK21 cells. And, Plaque forming assay of ZIKV and CVB3 was performed with Vero cells. Cells were seeded at a density of 2 × 10^5^ cells per well in a 24 well plate. 24 hours later, cell monolayers were washed and overlaid with 10-fold serial dilutions of culture supernatant from DENV and ZIKV infections. After one hour, the inoculum was replaced, cells were washed and overlaid with 1% CMC containing DMEM and 2% FBS. Cells were incubated at 37°C for 5 days for DENV and 4 days for ZIKV and the overlay media was removed and cells were fixed for 30 minutes with 3% formaldehyde. Fixed cells were stained for 15 minutes with 10% Crystal Violet and washed to visualize plaques. Plaques were counted, stated as plaque-forming unit (PFU/ml). EBOV titration was performed as previously described ^19^.

### Antibodies

Rabbit monoclonal antibody against TDRD3 was obtained from CST (5942S). Rabbit monoclonal antibody against TOP3B was obtained from ABcam (ab183520). Rabbit monoclonal antibodies against GFP were obtained from ThermoFisher Scientific (G10362). Mouse monoclonal antibodies against Actin were obtained from Santa Cruz Biotechnology (sc-47778). 4G2 anti-Flavivirus envelope Mouse monoclonal antibody was isolated from the DI-4G2-4-15 hybridoma, ATCC. Monoclonal Mouse Anti-Enterovirus (clone 5-D8/1) antibody was purchased from Agilent Dako (Code M7064).

### Plasmid construction

To construct CRISPR editing plasmids targeting *TDRD3* (pX330-TDRD gRNA1, pX330-TDRD gRNA1) or *TOP3B* gene (px330-TOP3B gRNA1, pX330-TDRD gRNA2), oligo sequences provided upon request, were purchased from IDT and were sub-cloned into pX330 as described previously ^20^. To generate pFLAG-TOP3B, human Full-length Top3B with FLAG-Tag sequence was amplified using the following set of primers: forward (5′-CCCAAGCTTATGGATTACAAGGATGACGACGATAAGAAGACTGTGCTCATGGTTGCT GAA-3′) and reverse (5′-CGCGGATCCTCATACAAAGTAGGCGGCCAGGGCTGACAT-3′). And the amplicon sequence was sub-cloned into pcDNA5/FRT/TO plasmid (Thermo Fisher Scientific). Plasmids GFP-tagged full length TDRD3 (pGFP-TDRD3), TOP3B (pGFP-TOP3B) and were described previously (Yang *et al*., 2010).

### Generation of TDRD3 and TOP3B knockout cells by the CRISPR/Cas9 system

To generate *TDRD3* KO cells, HuH-7 cells were co-transfected with pX330-TDRD3 gRNA1, pX330-TDRD3 gRNA2, and puromycin plasmid. At the second day of culture, puromycin (2mg/ml) was supplemented into the culture medium, and the cells with recombination of *TDRD3* and *PuroR* gene were selected for 3 days. Cells were subsequently seeded into 96-well plates at a density of 0.1 cells/well to generate single-cell clones. Only wells harboring a single clone were used. The gene KO was confirmed by Sanger sequencing of the *TDRD* gene and Western blotting as described below. HuH-7 *TOP3B* KO cells were generated in the same manner.

### Construction of tetracycline-inducible expression system

Stable tetracycline inducible HEK293 cell lines expressing FLAG-TOP3B were established with the Flp-in T-REx system (Thermo Fisher Scientific) by cotransfection of pFLAG-TOP3B and the pOG44 plasmid, which encodes Flp recombinase,. After transfection stable transfectants are selected using 100 μg/ml of hygromycin B and 15 μg/ml of blasticidin. Expression of the transgene was induced by the addition of 1 μg/mL doxycycline (DOX) in the culturing media.

### Immunoblotting

The HuH-7 wild-type (WT), HuH-7 *TDRD3* KO, or HuH-7 *TOP3B* KO cells were lysed in RIPA buffer (Cell Signaling Technologies) with 1X protease inhibitor cocktail (Cell Signaling Technologies) and centrifuged at 10,000 rpm for 3 min to pellet membranous cell debris. The cleared samples were separated by electrophoresis on a 4 to 12% Bis-Tris protein gels and transferred to nitrocellulose membranes. The membranes were blocked in blocking buffer for 1 h, followed by overnight incubation with primary antibody. The blots were then washed and incubated with anti-rabbit or anti-mouse fluorophore/DyLight FW 800-conjugated secondary antibody (Santa Cruz Biotechnology). Bands were visualized using Odyssey infrared scanner from Li-Cor.

### Virus growth assay in KO cells

5 × 10^4^ HuH-7 WT, TDRD3 KO or TOP3B KO cells were seeded on 24 well plates. After 24 h of culture, the cells were infected with DENV at an MOI of 0.2, ZIKV at an MOI of 0.2. The culture supernatants were collected at the 72 hours post-infection and the virus titer was measured by plaque forming assay. For YFV cells were infected at an MOI of 0.01, and for CVB3 at an MOI of 0.01. 72 hpi cells were stained with 4G2 mAb (YFV) or VP1 (CVB3) primary antibody, Alexa647-conjugated anti-mouse IgG mAb as a secondary antibody and DAPI, and the infection rate was obtained from the average of the fthree areas and shown as % infected cells. For CHIKV infections, cells were infected at an MOI of 0.05. 72 hpi cells were washed five times with PBS and then lysed with *Renilla* lysis buffer (Promega). Luciferase assays were performed using an EnSpire plate reader (PerkinElmer). For IAV infections, HuH-7 cell monolayers were washed once with DPBS, and IAV (strain Influenza A/Puerto Rico/8/1934 H1N1) was diluted in DPBS with 0.4% (w/v) BSA for infection at multiplicity of infection (MOI) of 0.01 PFU/cell, was incubated on cells at 37°C for 1 hr. Inoculations were removed, cells were washed twice with DPBS, overlaid with DMEM containing 0.4% (w/v) BSA, 0.1 μg/mL TPCK-treated trypsin, and incubated at 37°C.

Supernatants were collected for plaque assay at the indicated time points (24h, 48, 72h). Standard plaque assays were performed as previously described (PMID:20534532). In brief, for plaque assays, confluent monolayers of MDCK cells were washed once with DPBS. The collected supernatants containing virus were diluted in DPBS with 0.4% (w/v) BSA, and incubated on cells at 37°C for 1 hr. The medium was then removed, cells were washed twice with DPBS and replaced with MEM containing 0.6% oxoid agar, 1 μg/mL TPCK-treated trypsin and incubated at 37°C for 2 days. Cells were fixed in 3.7% (v/v) paraformaldehyde (PFA) in DPBS for 1 hr at room temperature, and stained with 1% (w/v) crystal violet, 20% (v/v) methanol in DPBS for 20 minOf the IAV experiment, the supernatant was collected over time (at 24, 48, 72 h.p.i). The virus titer was measured by plaque forming-assay.

### Immunofluorescence and imaging

IFA of cells infected with DENV, YFV or CVB3 was performed as described previously ^21^. Briefly, the cells were stained with 4G2 mAb as a primary antibody, Alexa647-conjugated anti-mouse IgG mAb as a secondary antibody and DAPI. VP1 antibody was used for the primary antibody against CVB3 infected cells. IFA images of the plates were taken with the 10× objective on an Opera Phenix High Content Screening System (PerkinElmer) and the images were analyzed using the associated Harmony® Office Software (PerkinElmer). Four microscopic areas per well were selected, and total and infected cells were counted based on Alexa647 and DAPI signal. The infection rate was obtained from the average of the four areas and shown as % infected cells.

### Transfections and Rescue experiment

5 × 10^4^ HuH-7 WT, TDRD3 KO or TOP3B KO cells were seeded on 24 well plates. After 24 h of culture, the cells were transfected with pGFP alone, or pGFP-TDRD3, or pGFP-Top3B and 24 hrs later transfected cells were split into a new plate and infected with DENV-2 at an MOI of 0.2. At 72 hr post-infection, viral titers in culture supernatants were measured by plaque-forming assay. Expression of exogenous GFP-tagged TDRD3 or TOP3B was confirmed by Western-blotting.

### Cross-liking and Immunoprecipitation (CLIP) of TOP3B

HEK293T-FLP-In Flag-TOP3B was seeded on a 10cm dish at the 2.2 × 10^6^ cells/plate and cultured over-night. The culture medium was exchanged to the medium with/without 1 μg/ml DOX to induce exogenous Flag-TOP3B, and at the same time, the cells were infected with DENV-2 at an MOI of 0.1. At the 48 h.p.i., the infected cells were crosslinked with 120 mJ UV light (wavelength 254 nm), or Mock-treated, UV: (−). The cells were lysed with lysis buffer (100mM Tris-HCl pH 7.4, 0.5 % NP-40, 0.5% Triton X-100, 100 mM NaCl_2_, 2.5 mM MgCl_2,_ and 0.1% sodium dodecyl sulfate (SDS)) and passed through 30G needle. After the centrifuge at 10,000 rpm for 15 min to pellet membranous cell debris, the sample was diluted into 0.25 mg/ml total protein and stored.

FLAG-TOP3 and crosslinked RNA complexes were immunoprecipitated from the 500 μl of total cell lysate by using Pierce ^TM^ Anti-DYKDDDK Magnetic Agarose beads (Thermo fisher scientific). After the extensive washing with high-salt buffer (1M NaCl_2_, 100mM Tris-HCl pH 7.4, 2.5m MgCl_2_, 0.5% NP-40, 0.5% Triton X-100, 0.1% SDS), the input cell-lysate, the depleted supernatant after IP and 25% of the beads were analyzed with Western-blotting and induced expression and IP were confirmed.

To degradate crosslinked proteins and release the precipitated RNA, the input cell-lysate and remained 75% of the beads were treated with Proteinase K (NEB) for 15 min at 37 °C. RNA was extracted with Direct-zol RNA kit (Zymo Research), reverse transcription was conducted with random primers using the High Capacity cDNA Reverse Transcription Kit (Thermo Fisher Scientific). EEF1A1 RNA (forward 5’-TTGGTTCAAGGGATGGAAAG-3’, reverse 5’-TTGTCAGTTGGACGAGTTGG-3’), CELSR2 RNA (forward 5’-CTCTCTGCACAGCACCAG-3’, reverse 5’-CCTTGGCCAGGGTTCAG-3’) or DENV-2 genomic RNA (forward 5’-GAAATGGGTGCCAACTTCAAGGGT-3’, reverse 5-GAAATGGGTGCCAACTTCAAGGGT-3’) was detected and measured by real-time quantitative PCR (RT-qPCR) using PowerUP SYBR Green Master Mix (Thermo Fisher Scientific) in StepOnePlus Real-Time PCR System (Thermo Fisher Scientific). Primers used in RT-qPCR are shown in Table S1. Co-precipitation of the RNA was shown as % Total RNA and Fold Enrichment against DOX (−) condition.

### Statistical analysis

In all figures, error bars represent the standard deviation (S.D.). Asterisks denote level of statistical significance: * p < 0.05, ** p < 0.01, *** p<0.001 and **** p < 0.0001. To ensure the normal distribution of the value, the virus titer was statistically analyzed after the Log_10_ transfer. For the comparison of virus titer between HuH-7 WT and TDRD3 or TOP3B KO cells, an unpaired student’s t-test was performed. This analysis was performed at each time point in the IAV growth assay. For the side by side comparison of virus titer in HuH-7 WT, TDRD3 KO and TOP3B KO cells and rescue assay of viral replication by exogenous expression of TDRD3 or TOP3B, Tukey’s multiple comparison test was performed after the Ordinary One-way ANOVA. In the % Total RNA of the CLIP experiment, Dunnett’s multiple comparison test against EEF1A1 was performed after the Ordinary One-way in each experimental condition. In the comparison of fold enrichment of DENV-2 genomic RNA of the CLIP experiment, Tukey’s multiple comparison test was performed after the Ordinary One-way ANOVA. Statistical analyses were conducted in Prism8 (GraphPad Software).

